# Endogenous Tenocyte Activation Underlies the Regenerative Capacity of Adult Zebrafish Tendon

**DOI:** 10.1101/2023.02.04.527141

**Authors:** Stephanie L. Tsai, Steffany Villasenor, Rishita Shah, Jenna L. Galloway

## Abstract

Tendons are essential, frequently injured connective tissues that transmit forces from muscle to bone. Their unique highly ordered, matrix-rich structure is critical for proper function. While adult mammalian tendons heal after acute injuries, endogenous tendon cells, or tenocytes, fail to respond appropriately, resulting in the formation of disorganized fibrovascular scar tissue with impaired function and increased propensity for re-injury. Here, we show that unlike mammals, adult zebrafish tenocytes activate upon injury and fully regenerate the tendon. Using a full tear injury model in the adult zebrafish craniofacial tendon, we defined the hallmark stages and cellular basis of tendon regeneration through multiphoton imaging, lineage tracing, and transmission electron microscopy approaches. Remarkably, we observe that the zebrafish tendon can regenerate and restore normal collagen matrix ultrastructure by 6 months post-injury (mpi). We show that tendon regeneration progresses in three main phases: inflammation within 24 hours post-injury (hpi), cellular proliferation and formation of a cellular bridge between the severed tendon ends at 3-5 days post-injury (dpi), and re-differentiation and matrix remodeling beginning from 5 dpi to 6 mpi. Importantly, we demonstrate that pre-existing tenocytes are the main cellular source of regeneration. Collectively, our work debuts the zebrafish tendon as one of the only reported adult tendon regenerative models and positions it as an invaluable comparative system to identify regenerative mechanisms that may inspire new tendon injury treatments in the clinic.

## Introduction

Tendons are highly specialized structures that connect and transmit forces from muscle to bone, enabling movement. Compared to other tissues, tendons are relatively hypocellular. The mass of the tendon is predominantly comprised of a dense extracellular matrix composed of highly aligned type I collagen fibers running uniaxially from the myotendinous junction (MTJ) through the midbody to the tendon-bone attachment, or enthesis. Deeply embedded within the matrix are organized arrays of tendon cells, or tenocytes, which are elongated, stellate-shaped cells with long processes that form an intricate network with neighboring cells via gap junctions^1–3^. While tendons can vary in size and morphology, the highly aligned matrix and tenocyte network are stable features universally required for efficient force transmission and mechanotransduction along the length of the tendon^4–6^.

As tendons play a pivotal role in everyday movement, they are frequently injured. Unfortunately, the unique, highly ordered tissue architecture cannot be restored in mammals post-injury. Severe tendon injuries including full tears result in the formation of disorganized fibrovascular scar tissue with impaired function and a higher likelihood of both re-injury as well as developing joint degenerative conditions^7^. Although injuries to tendons and other joint connective tissues are estimated to account for 45% of all musculoskeletal injuries, effective treatments are limited and often result in surgical intervention, which is costly and variable in success^8–10^. Better therapeutic strategies to treat tendon injuries are needed in the clinic to improve patient quality of life as well as ameliorate a mounting healthcare burden.

To this end, most research efforts have been largely focused on understanding the mechanisms underlying adult mammalian fibrotic tendon healing to identify new therapeutic strategies to augment existing clinical treatments. Yet, important lessons on how to achieve tendon regeneration rather than scarring can be learned from natural examples that have been recently reported including in neonatal mice^11–13^ and larval zebrafish^14^. These tendon regenerative models can serve as powerful systems to identify mechanisms required for tendon regeneration which may ultimately accelerate the advancement of clinical strategies. However, the previous studies in zebrafish and mice are examples of regeneration set during developmental stages in which the tendon is still actively growing and maturing^15,16^. As cellular plasticity is higher during tissue formation, the mechanisms directing neonatal or larval regeneration may differ between development and adulthood. An adult tendon regenerative model would therefore be invaluable to the field as a comparative paradigm to understand mechanisms driving proper regeneration.

While mammalian regenerative capacity declines from embryonic and postnatal stages to adulthood, zebrafish retain the remarkable ability to regenerate various tissues and organs as adults including their heart and spinal cord^17^; however, tendon regeneration has yet to be examined. Here, we demonstrate that adult zebrafish regenerate their tendon following full transection. We delineate hallmark processes of tendon regeneration and reveal that the pre-existing tenocytes are the main cellular source of regenerated tendon tissue.

## Results

### The adult zebrafish tendon can regenerate after acute injury

To determine if adult zebrafish tendons regenerate, we performed full transection injuries on the craniofacial maxillary superficial tendon (MST) in *scxa:mCherry* zebrafish and monitored the recovery (Fig. 1a-d). The MST connects the maxilla to one of the jaw adductor muscles, specifically classified as the A0 adductor muscle (Fig. 1a-b)^18,19^. The midbody of the MST contains a short segment with only midbody tenocytes and a longer segment which runs along the adductor muscle and contains a mixture of both MTJ cells and midbody tenocytes. In the latter region, all cells express the tendon marker *tnmd*, and cells only in direct contact with the muscle express the myotendinous marker *col22a1* (Fig. 1c, Supplementary Fig. 1)^20–23^. For technical reproducibility, we chose to perform a full transection of the tendon at its midpoint towards the beginning of the longer segment. We also strategically chose the MST because its superficial nature makes it ideal for experimental use as it is easy to access, surgically manipulate, and image. In addition, while it functions in closing the mouth^19^, it is not required for feeding allowing fish to eat normally and survive following surgery.

**Figure 1.**
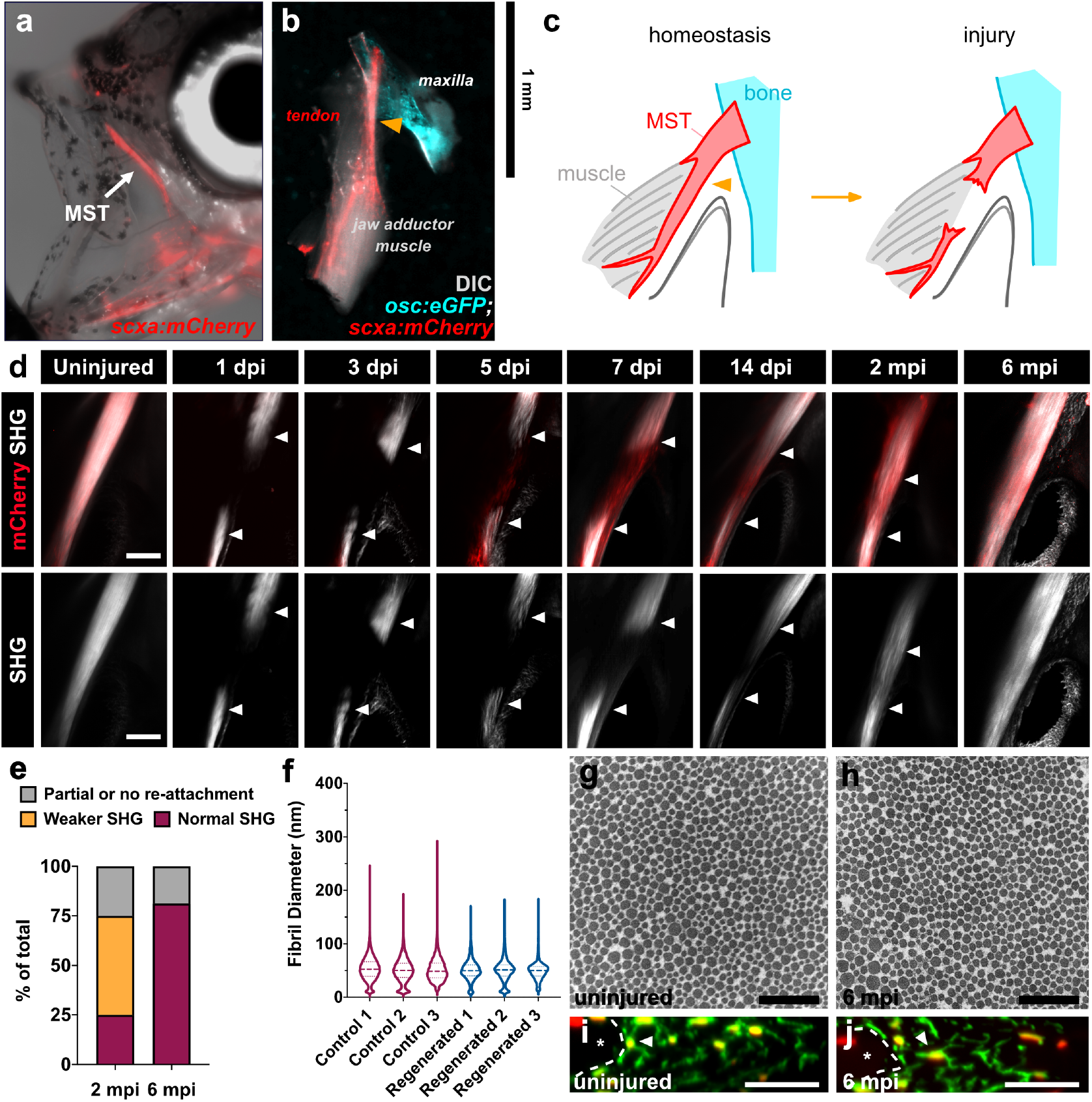
The adult zebrafish can fully regenerate after acute injury. (a) Epifluorescence image of an adult *scxa:mCherry* zebrafish with a brightfield overlay to demonstrate the position of the maxillary superficial tendon (MST) (denoted by white arrow). (b) Epifluorescence image of a normal uninjured MST musculoskeletal circuit (i.e. maxilla, MST, jaw adductor muscle (A0)) dissected from an *osc:eGFP;scxa:mCherry* adult zebrafish with a DIC overlay. Orange arrowhead denotes where the injury is made. (c) Graphical schematic of the MST before and after injury. (d) 2-photon images of the MST at various timepoints after injury in *scxa:mCherry* zebrafish with SHG signal both overlaid and shown separately. White arrowheads denote the severed tendon ends. Scale bar, 100 μm. (e) Stacked bar graph showing the breakdown of zebrafish exhibiting partial or no reattachment, weaker SHG signal, or fully restored SHG signal at 2- and 6-months post-injury (mpi). (f) Violin plot illustrating the collagen fibril diameter distribution in 3 age-matched individual uninjured and regenerated MSTs at 6 mpi. Means of uninjured controls 1, 2, and 3 were 53.33, 50.24, and 51.35 nm, respectively. Means of injured MSTs 1, 2, and 3 were 50.62, 50.31, and 49.41 nm respectively. (g-h) Representative 50,000x TEM micrographs from age-matched uninjured (g) and regenerated tendons at 6 mpi (h). Scale bar, 500 nm. (i-j) Representative images of a cross-sectional re-slice view of 2-photon z-stacks from anti-mCherry stained (shown in green) *scxa:mCherry* MSTs from age-matched control uninjured (i) and regenerated MSTs at 6 mpi (j). White dotted lines outline muscle boundary. Scale bar, 10 μm.

We performed 2-photon imaging at different timepoints post-injury to monitor the recovery, which allowed us to simultaneously visualize *scxa:mCherry* expression and second harmonic generation (SHG) signal to assess collagen fiber alignment (Fig. 1d). The uninjured tendon exhibits both strong *scxa:mCherry* expression and SHG signal as expected. At 1 and 3 days post-injury (dpi), there is a loss of *scxa:mCherry* expression and SHG signal at the injury site. However by 5 and 7 dpi, a *scxa:mCherry+* bridge between the severed tendon ends can be observed and appears to become more organized by 14 dpi. By 2 months post-injury (mpi), SHG signal begins to return at the injury site and appears to be fully restored by 6 mpi. When we examined the qualitative breakdown of SHG signal restoration following injury, we observed that ~25% of injured fish (N=3/12) had fully restored SHG signal at the injury site at 2 mpi whereas ~50% (N=6/12) displayed weaker SHG signal (Fig. 1e). By 6 mpi, ~81.25% of injured fish (N=13/16) exhibited a full restoration of SHG signal, indicating collagen fiber alignment in the tendon matrix had returned. We also observed that some fish exhibited partial or no re-attachment of the severed tendon ends (N=3/16), leading to a complete failure of SHG signal restoration.

We next sought to examine whether the collagen matrix ultrastructure and tenocyte morphology/network regenerated following injury. To assess if the collagen fibril diameter distribution was re-established, we performed transmission electron microscopy (TEM) of control and injured tendons at 6 mpi (Fig. 1f-h). We observed no significant difference in collagen fibril diameter distributions between control and injured tendons at 6 mpi, suggesting that the collagen matrix was fully restored. Furthermore, tenocytes in the injury site reassumed a normal elongated tenocyte morphology with thin processes extending between neighboring cells, indicating the tenocyte network is re-established in injured tendons by 6 mpi (Fig. 1i-j). Altogether, these data strongly show that the adult zebrafish tendon fully regenerates following acute injury, unlike their mammalian counterparts.

### Tendon injury triggers a swift innate immune response, cellular proliferation, and collagen fiber deposition during the first week post-injury

As the formation of an organized *Scx+* cellular bridge does not occur following full transection in mammalian tendons, we sought to better characterize early events during zebrafish regeneration. Masson’s trichrome staining of the regenerating tendon at 1, 3, and 5 dpi revealed a rapid sequence of events surrounding bridge formation during the first week post-injury. At 1 dpi, there is a heavy infiltration of cells, many of which exhibit monocytic and granulocytic morphologies (e.g. rounded, large multi-lobed nuclei), suggesting the onset of an innate immune response. By 3 dpi, a cellular bridge has begun to form between the two severed tendon ends containing cells that exhibit fibroblastic morphologies. At 5 dpi, the beginnings of collagen deposition are evident as seen by the presence of collagen fibers stained in blue (Fig. 2).

**Figure 2.**
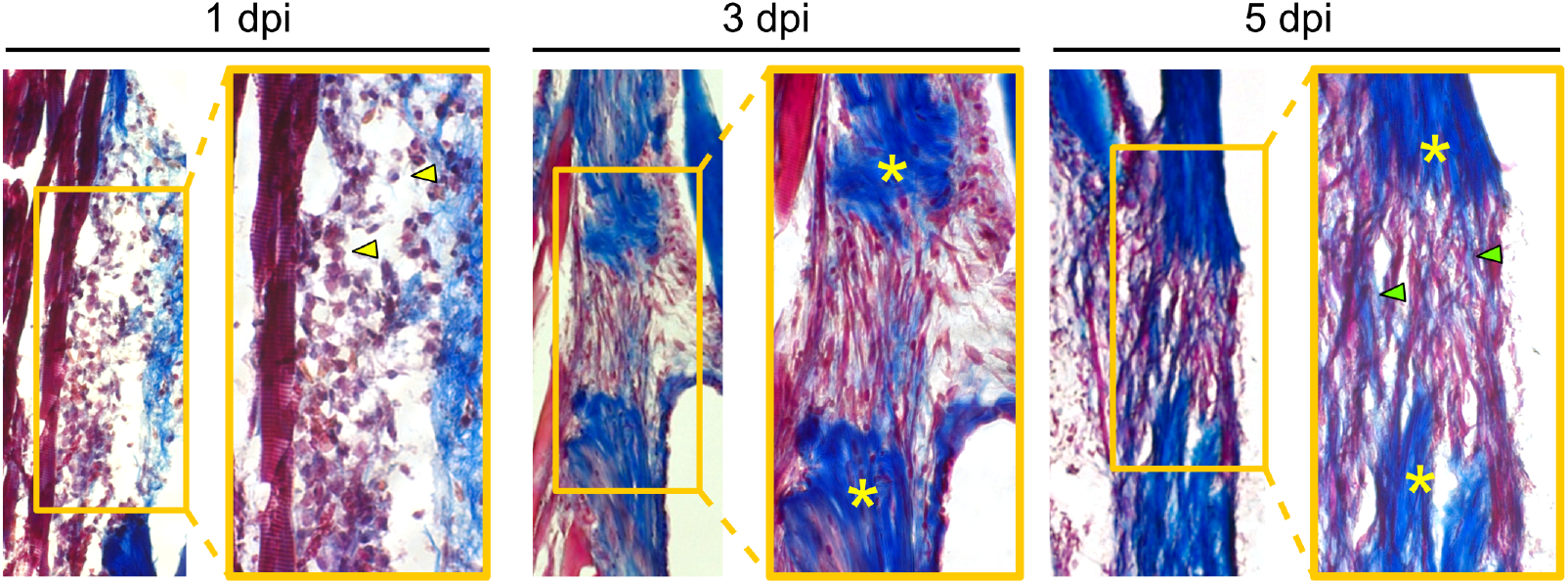
Tendon regeneration proceeds through a rapid series of phases within the first week post-injury. Masson’s trichrome staining of sections from regenerating tendons at 1 (left), 3 (middle), and 5 (right) dpi. Heavy infiltration of cells with myeloid-like morphologies can be seen at 1 dpi (yellow arrowheads). At 3 dpi, a fibroblastic bridge connecting the two severed tendon ends is evident. By as early as 5 dpi, the beginnings of collagen matrix deposition into the injury site are observed (green arrowheads). Yellow asterisks denote severed tendon ends and images taken at 10x magnification. Dpi, days post-injury.

To examine innate immune cell dynamics during tendon regeneration in more depth, we performed time course 2-photon imaging of *Tg(mpx:eGFP)* and *Tg(mpeg:eGFP)* lines to examine neutrophil and macrophage infiltration, respectively (Fig. 3a-b). Overall, both cell types demonstrated significant dynamic changes in infiltration that were relatively similar after injury (****p<0.0001, one-way ANOVA). We observed little to no neutrophils or macrophages in the homeostatic tendon at 0 dpi. However, by 12 hours post-injury (hpi), there was a rapid and significant increase in both cell types at the site of injury (*mpx+* cells: 0.21% (0 dpi) vs. 15.09% (12 hpi), ****p<0.0001, *mpeg+* cells: 0.0% (0 dpi) vs. 11.78% (12 hpi), ****p<0.0001). By 1 dpi, the percentage of neutrophils in the injury site significantly decreased (15.09% (12 hpi) vs. 6.34% (1 dpi), ****p<0.0001) whereas the percentage of macrophages did not significantly change from 12 hpi. However, the percentage of both immune cell types continued to decrease over time from 1 to 5 dpi, at which point near basal levels were reached (*mpx+* cells: 6.34% (1 dpi) vs. 0.91% (5 dpi), **p<0.01, *mpeg+* cells: 8.20% (1 dpi) vs. 1.26% (5 dpi), *p<0.05). At 14 dpi, virtually no neutrophils or macrophages remained at the injury site. These data indicate that tendon injury triggers a robust yet transient innate immune response that peaks at 12 hpi and begins to decline thereafter to basal levels at 5 dpi.

**Figure 3.**
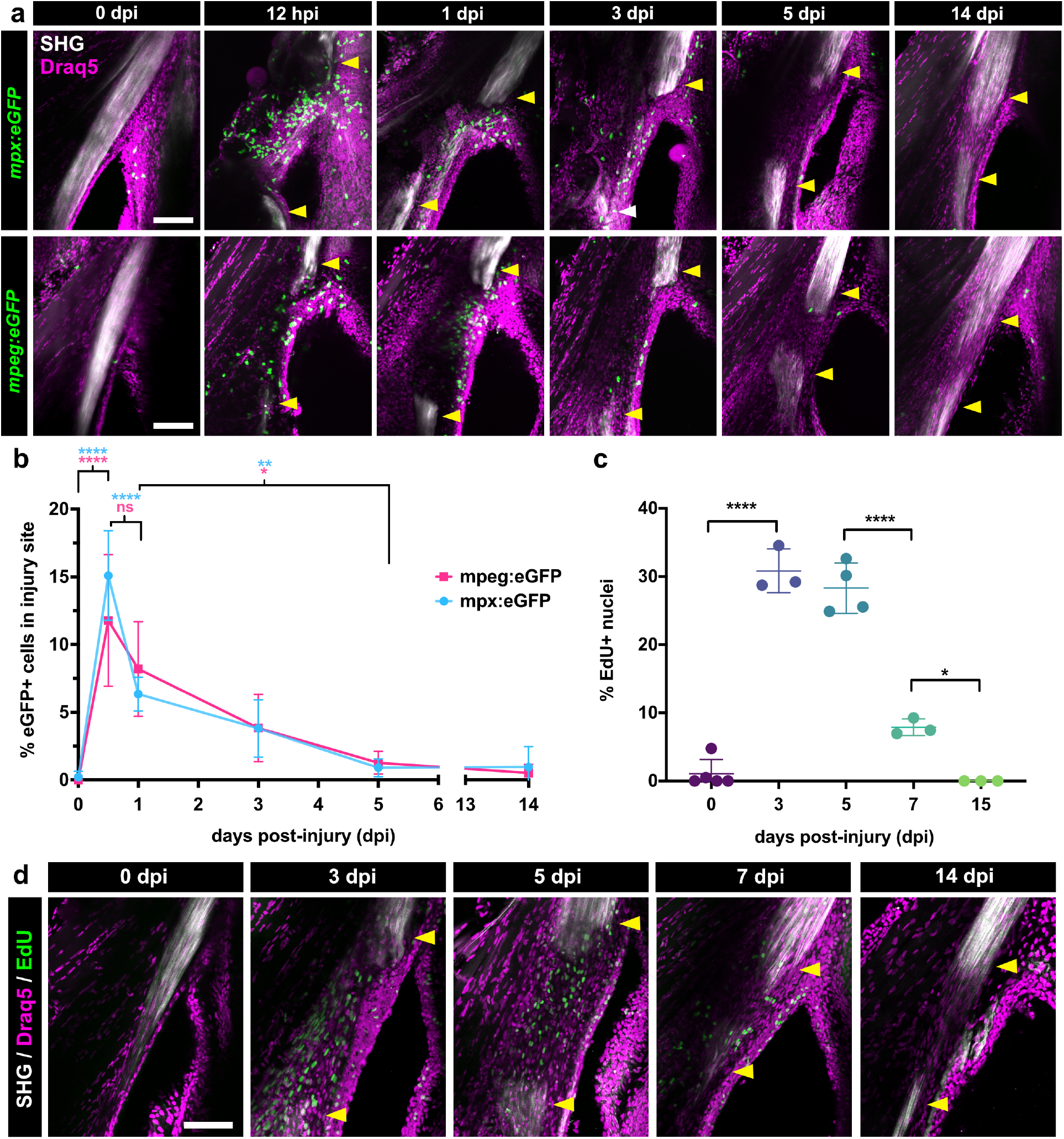
Tendon injury triggers a rapid innate immune response followed by a wave of cellular proliferation. (a) 2-photon imaging of regenerating tendons from *mpx:eGFP* (top row) or *mpeg1:eGFP* (bottom row) zebrafish at different timepoints post-injury to examine neutrophil and macrophage dynamics, respectively. SHG signal is overlaid along with a Draq5 nuclear counterstain. Yellow arrowheads denote severed tendon ends. Scale bar, 100 μm. (b) Quantification of the percentage of *mpx:eGFP+* and *mpeg1:eGFP+* cells out of total Draq5+ cells in the injury site during the first 2 weeks post-injury. One-way ANOVA analysis was employed for statistical analysis with Tukey’s multiple comparison tests between different time points. Sample sizes were as follows: *mpx:eGFP* – 0, 1, 14 dpi: N=4; 0.5, 3, 5 dpi: N=5; *mpeg1:eGFP* – 0.5 dpi: N=6; 0, 1, 3 dpi: N=5; 5 dpi: N=4; 14 dpi: N=3. ****p<0.0001, **p<0.01, *p<0.05. (c) Quantification of the percentage of EdU+ cells out of total Draq5+ cells in the injured area at 0, 3, 5, 7, and 15 dpi. One-way ANOVA analysis was employed for statistical analysis with Tukey’s multiple comparison tests between timepoints. Sample sizes were as follows: 0 dpi: N=5; 3, 7, 15 dpi: N=3; 5 dpi: N=4. ****p<0.0001, *p<0.05. (d) Representative 2-photon time course imaging of EdU+ cells (in green) at different time points post-injury. SHG signal is overlaid with Draq5 nuclear staining. Yellow arrowheads denote severed tendon ends. Scale bar, 100 μm.

We next asked whether bridge formation coincides with a peak in cellular proliferation of a potential progenitor population that ultimately regenerates the tendon. To examine proliferating cells, we pulsed regenerating zebrafish with EdU 24 hours prior to imaging to assess the percentage of proliferating cells at different time points post-injury (Fig. 3c-d). We observed significant changes in the percentage of EdU+ cells during early stages of regeneration (****p<0.0001, one-way ANOVA). As expected, little to no EdU+ cells were present in the uninjured tendon at 0 dpi. However, there was a statistically significant increase in the percentage of EdU+ cells at the injury site at 3 dpi (1.05% (0 dpi) vs. 30.82% (3 dpi), ****p<0.0001) which was steadily maintained through 5 dpi, began to significantly decrease at 7 dpi (28.30% (5 dpi) vs. 7.88% (7 dpi),****p<0.0001), and ultimately declined down to basal levels at 15 dpi (7.88% (7 dpi) vs. 0.00% (15 dpi), *p<0.05). Collectively, these data indicate that tendon injury triggers a series of hallmark events within the first 5 days after injury including a strong innate immune response followed by increased cellular proliferation and the onset of collagen matrix deposition.

### Endogenous tenocytes are a major cellular source of tendon regeneration

While neonatal mammalian *Scx*-lineage tendon cells can respond to injury and regenerate the Achilles tendon following full transection, adult *Scx*+ tenocytes fail to respond upon the same injury and do not contribute to healing^12^. Therefore, we asked whether adult zebrafish tenocytes may differ from their counterparts in mice and retain the ability to respond to injury and contribute to tendon regeneration. To examine this question, we generated a *scxa:creERT2* BAC transgenic zebrafish line to perform lineage tracing of tenocytes during regeneration (Fig. 4). We first validated the specificity of the *Tg(scxa:creERT2*) line in larvae by performing both double *in situ* hybridization chain reaction (HCR) of *scxa/cre* and 4-hydroxy-tamoxifen (4-OHT) labeling during tendon development. HCR double *in situ* hybridization of *scxa* and *cre* expression in *scxa:creERT2* larvae at 4 days post-fertilization (dpf) confirmed *cre* expression in *scxa+* craniofacial tendons and ligaments including the hyohyal (hh) and sternohyoidus (sh) tendons as well as the lateral ligaments (l) (Fig. 4a). Furthermore, 4-OHT induction of *scxa:creERT2;ubi:zebrabow* larvae from 48-96 hours post-fertilization (hpf) efficiently and specifically labeled cells within tendons in the craniofacial region including the sh tendon as well as the myosepta in the trunk region (Fig. 4b-d). Finally, we performed 4-OHT labeling in adult *scxa:creERT2;ubi:zebrabow* zebrafish and observed labeling of ~40-60% of tenocytes in the MST during homeostasis (Fig. 4e, Supplementary Fig. 2). Importantly, we observed that the percentage of CFP+/YFP+ labeled tenocytes remained stable after 7 days post-induction, thus for all subsequent adult lineage tracing experiments we chose to perform the injuries at 7 days post 4OH-T administration (Supplementary Fig. 2). Altogether, these data demonstrated the specificity and utility of this transgenic line for tendon lineage tracing studies.

**Figure 4.**
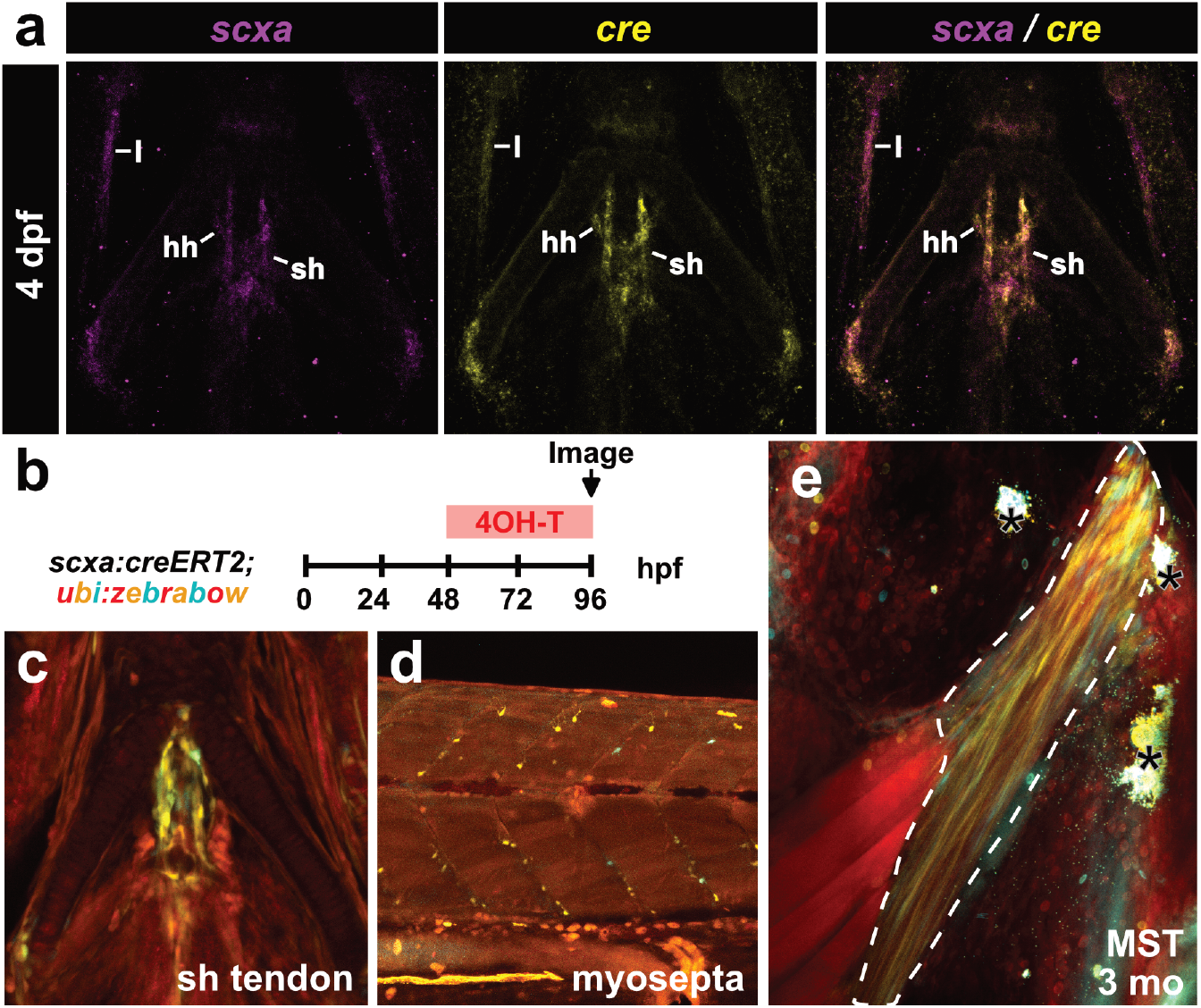
Generation and validation of a *scxa:creERT2* transgenic line. (a) HCR double *in situ* hybridization of *scxa* and *cre* at 4 days post-fertilization (dpf) shows overlap in expression in craniofacial tendons and ligaments. l, lateral ligament; sh, sternohyoidus tendon; hh, hyohyal tendon. Scale bar, 100 μm. (b) Schematic of 4OH-T labeling experiment to test tendon labeling in *scxa:creERT2; ubi:zebrabow* larvae. (c-d) Representative confocal images from the 4OH-T labeling validation experiment in b. The labeled SH tendon is shown in c and labeled myoseptal cells in the trunk region are shown in d at 96 hours post-fertilization (hpf). Scale bar, 100 μm. (e) Representative 2-photon image of 4OH-T labeling in the 3 month-old adult MST (outlined in white dotted line). Asterisks denote autofluorescent pigment in the skin. Scale bar, 100 μm.

To determine if *scxa*+ tenocytes contribute to tendon regeneration, we induced CFP and/or YFP labeling in *scxa:creERT2;ubi:zebrabow* zebrafish through 4-OHT administration one week prior to injury and examined the contribution of *scxa*-lineage cells to the regenerating tendon at 4 and 14 dpi (Fig 5a). At 4 dpi, we observed the presence of CFP+ and/or YFP+ *scxa*-lineage cells at the injury site that appeared to have a more rounded, less elongated morphology (Fig. 5a-b). In addition, CFP- and YFP-labeled cells were co-labeled with EdU in both of the severed tendon ends and at the injury site at 4 dpi (Fig. 5c-d), demonstrating that pre-existing tenocytes proliferate and migrate to the site of injury. By 14 dpi, CFP- and YFP-labeled cells re-adopted an elongated morphology at the injury site, indicating reacquisition of mature tenocyte features (Fig. 5b). Therefore, our data indicates that pre-existing tenocytes proliferate, migrate to the wound, and regenerate the tendon, collectively positioning them as the main cell source of adult zebrafish tendon regeneration.

**Figure 5.**
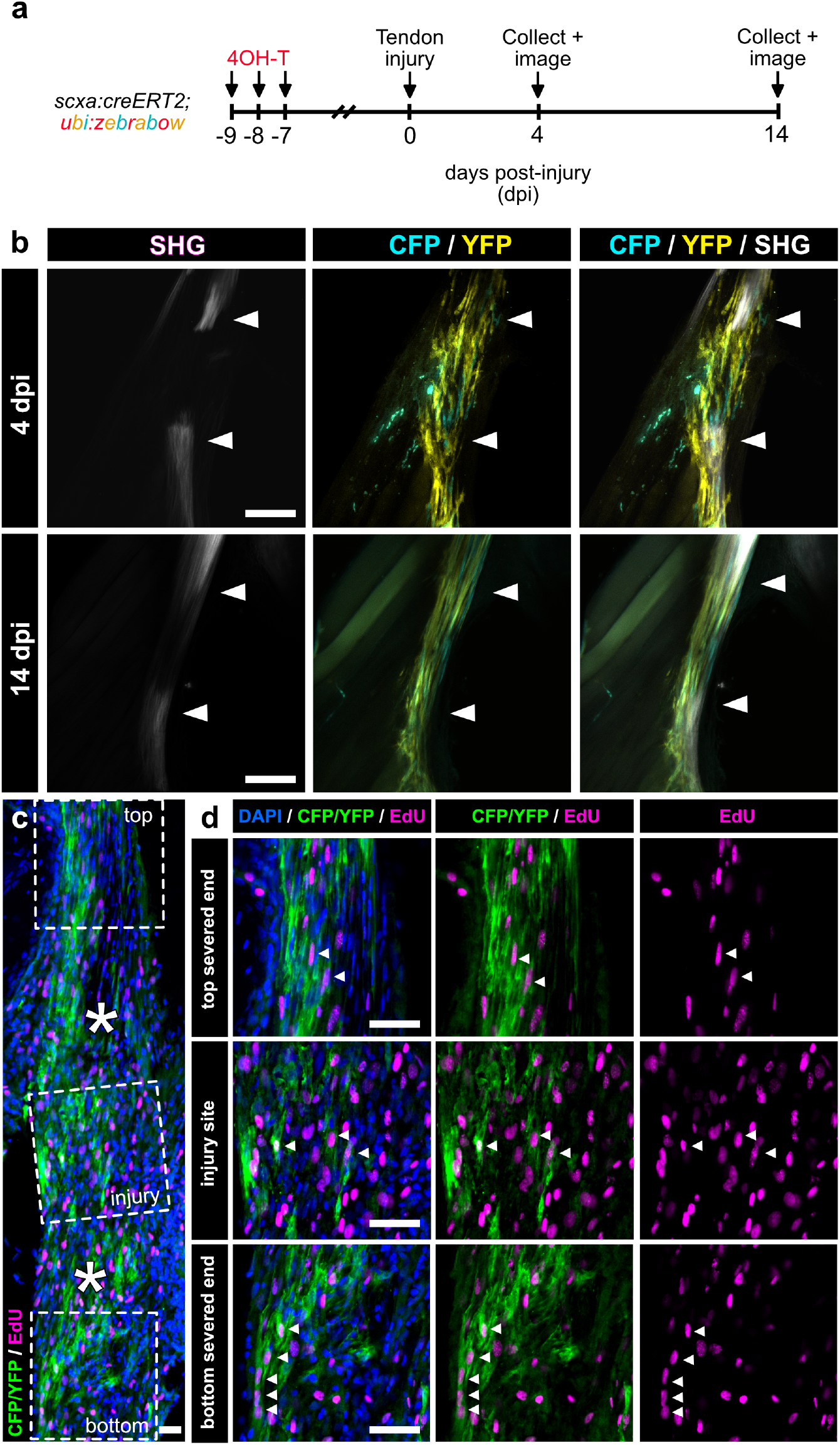
Pre-existing tenocytes are a major cell source of tendon regeneration. (a) Experimental schematic of *scxa:creERT2; zebrabow* lineage tracing experiment during tendon regeneration. (b) 2-photon imaging showing CFP+ and YFP+ cells infiltrating the injury site in *scxa*-lineage traced regenerating tendons at both 4 and 14 days post-injury (dpi). SHG signal is shown either separate or overlaid. White arrowheads denote severed tendon ends. Scale bar, 100 μm. (c) Representative confocal image of an anti-CFP/YFP stained (in green) section of a regenerating tendon at 4 dpi coupled with EdU labeling (in magenta). Higher magnification of regions of the top severed end, injury site, and bottom severed end are shown in d. Asterisks denote the severed tendon ends. Scale bar, 25 μm. (d) Higher magnification confocal imaging of CFP/YFP stained cells co-labeled with EdU in the top and bottom severed tendon ends as well as the site of injury. White arrowheads denote examples of CFP/YFP+ cells co-labeled with EdU. Scale bar, 25 μm.

## Discussion

Our work presents one of the first natural adult models of tendon regeneration and sets the groundwork for utilizing the zebrafish tendon as a means for uncovering molecular mechanisms required for proper restoration. We demonstrate that the adult zebrafish tendon can regenerate following an acute full transection injury and identify endogenous tenocytes as the main cellular source of regeneration. Tendon regeneration progresses through three main phases: inflammation, tendon bridge formation via tenocyte proliferation and migration, and re-differentiation/maturation coupled with matrix remodeling (Fig. 6). These stages advance in a semi-overlapping sequence within the first week post-injury, with the exception of tenocyte differentiation/maturation and matrix remodeling which spans up to 6 mpi, altogether demonstrating the immense time required to rebuild the distinct collagen matrix ultrastructure.

**Figure 6.**
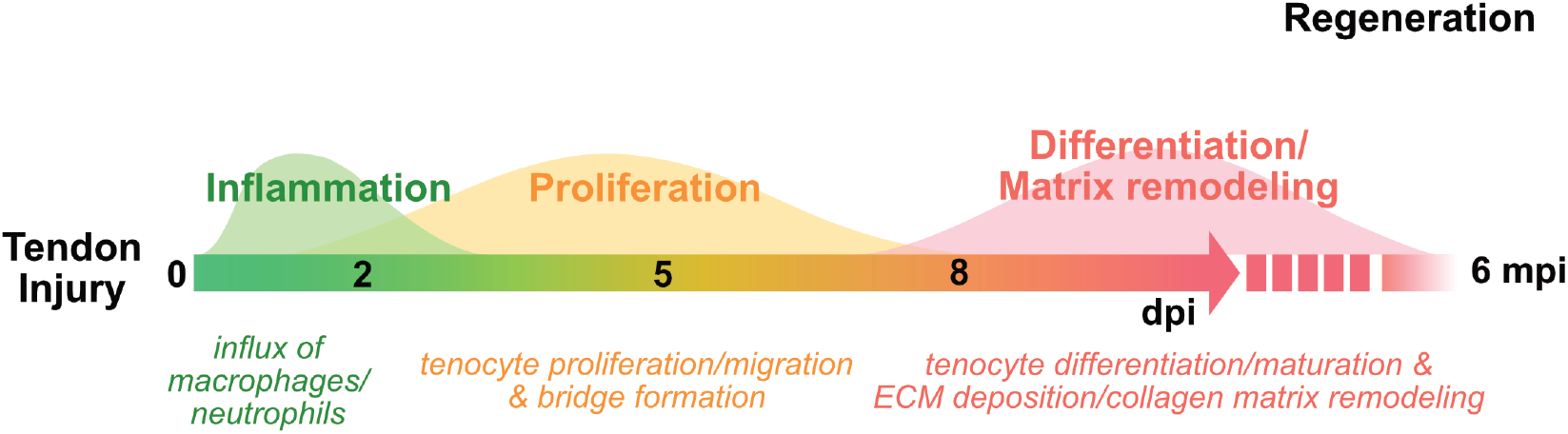
Hallmarks of adult zebrafish tendon regeneration. Schematic detailing the timeline of key processes following acute tendon injury and regeneration. dpi, days post-injury; mpi, months post-injury.

The inherent regenerative capability of the adult zebrafish tendon may be directly linked to the ability of their tenocytes to proliferate and migrate to the site of injury, forming a regenerative bridging tissue between the two severed tendon ends. Adult mammalian tenocytes appear to lack this ability as it has been shown that following full transection of the mouse Achilles tendon, *Scx*+ tenocytes in the severed ends incorporate EdU similar to our observations, but fail to infiltrate the site of injury^12^. Instead, the authors observed the recruitment and persistence of αSMA-expressing cells, which eventually led to the formation of fibrotic scar tissue. With this context in mind, our work pinpoints clear differences in cellular and molecular mechanisms leading to tendon regeneration in the zebrafish and fibrotic healing in mice following acute injuries. Foremost, it positions the adult zebrafish tendon as an informative comparative model system to elucidate both instructive and inhibitory cues required for driving the initiation of tenocyte activation in regeneration and lack thereof in mice.

While our work identifies the cellular basis of tendon regeneration in the adult zebrafish, many questions remain as to the extent of tenocyte plasticity during regeneration. We show that the tenocyte bridge begins to form by as early as 3 dpi and is initially *scxa:mCherry*-negative, but gradually gains *scxa:mCherry* expression over time. By 7 dpi, the tenocyte bridge is largely *scxa:mCherry*+. Labeled tenocytes and/or their descendants which proliferate and migrate to the injury site at 4 dpi are more rounded and eventually re-adopt an elongated morphology, a key characteristic associated with tenocyte maturation, during later stages. One interpretation of these pieces of evidence suggests that tenocytes revert to a more immature cell state during the initial stages of regeneration. However, whether they dedifferentiate to a multipotent progenitor that can give rise to other connective tissues including cartilage or to a lineage-restricted tendon progenitor remains to be determined.

Differentiating between these possibilities will likely require a deeper understanding of gene regulatory networks (GRNs) regulating tendon progenitor specification, as *Scx* remains the earliest reported tendon marker^24,25^. Tendons in the craniofacial region like the MST are derived from cranial neural crest cells (CNCCs), while those in limbs are derived from the lateral plate mesoderm^26–28^. Regional differences in the GRNs underlying tendon development and regeneration are therefore likely to exist and have yet to be fully explored. Whether the distinct developmental origins may influence the mechanisms driving regeneration and/or overall regenerative capacity remains an open and interesting question. Future comparative studies examining the regenerative ability across non-craniofacial tendons in the zebrafish will be instrumental in extracting basic molecular principles that are commonly required for tendon regeneration regardless of differences in anatomical location or developmental origin. Notably, deciphering how precise tendon lineage specification and maintenance is ensured upon injury and regeneration can provide insight into targetable mechanisms that may be exploited clinically to prevent heterotopic ossification of tenocytes, which commonly occurs after mammalian tendon injury and compromises tissue function^29^.

Importantly, it remains unclear whether all *scxa+* tenocytes in the adult zebrafish tendon are equally competent to respond to injury. It is possible that specific subpopulations of tenocytes are more (or less) poised to respond to injury and regenerate the tendon. In mammals, several adult tendon stem/progenitor populations have been identified *in vivo* which contribute to tendon healing^30–32^. Determining whether the zebrafish tendon contains similar subpopulations and pinpointing mechanisms that may lead to differential responses during regeneration versus fibrosis will be important next steps towards understanding the pathways and cell types that are advantageous or refractory for regeneration. Notably, it is possible that in addition to tenocytes, other neighboring cell types including interstitial connective tissues may also contribute and/or play a role. As our molecular understanding of the surrounding interstitial tissues is currently limited, further characterization of the MST and its neighboring tissues using single cell -omics approaches will facilitate the development of new genetic tools to answer these outstanding questions.

In all, our work debuts the adult zebrafish as a powerful genetic model which can be utilized to construct a blueprint of molecular and cellular mechanisms required for proper tendon regeneration. In combination with existing mammalian tendon healing models, our findings open up an invaluable opportunity to utilize comparative cross-species genomics approaches to enable the identification of genetic regulatory and signaling dynamics essential for driving regeneration versus fibrotic healing. Identifying these mechanisms will be instrumental in accelerating innovation of effective tendon injury treatments in the clinic.

## Materials and Methods

### Animal husbandry and zebrafish lines

All zebrafish were housed and maintained according to the MGH Institute for Animal Care and Use Committee (IACUC) protocol guidelines (Protocol #: 2012N000167). For the experimental data generated in this study, we utilized the following zebrafish transgenic lines: *TgBAC(scxa:mcherry)^33^, Tg(mpx:eGFP)^34^, Tg(mpeg:eGFP)^35^, Tg(ubi:zebrabow)^36^, and TgBAC(scxa:creERT2)*.

### Tendon injuries

All surgeries were performed according to MGH IACUC protocol guidelines (Protocol #: 2012N000167). Adult zebrafish from 6-15 months old were utilized. To perform adult tendon injuries, zebrafish were anesthetized in 0.015% tricaine and gently pinned down with staples on to an agarose plate on their side to immobilize them. Using a small pair of dissecting scissors, the craniofacial maxillary superficial tendon (MST) was fully transected at its midpoint. The zebrafish was then returned into normal system water and monitored daily for recovery.

### Transmission electron microscopy (TEM) analysis of the zebrafish tendon

TEM of uninjured control and regenerated tendons (6 months post-injury) was performed at the Shriners Hospitals for Children in Portland, Oregon. A total of 3 control and 3 regenerated tendons were analyzed. Collagen fibril diameters from each sample were measured using Fiji. For each individual sample, 5-10 50,000x images across 3-4 different planes were blinded and analyzed through the injury site or the corresponding uninjured area in the control tendons. A total of ~6000-35000 fibrils were measured for each sample.

### 2-Photon microscopy

For the time course imaging of *scxa:mCherry* fish, adults were euthanized in 0.1% tricaine in order to immobilize the heart and pinned down onto an agarose plate with the mouth opened to allow for full extension of the tendon. The MST was imaged at 25x magnification at various time points pre- and post-injury. To visualize neutrophils and macrophages, *mpx:eGFP* and *mpeg:eGFP* fish were euthanized and the heads were fixed overnight in 4% paraformaldehyde at 4 degrees. The tendons were then dissected out of the heads and stained overnight in Draq5 in 1% Triton-X/D-PBS. The stained tendons were embedded into 1% low melting point (LMP) agarose and imaged on the 2-photon. For all 2-photon imaging, second harmonic generation signal was acquired and overlaid to visualize the tendon collagen matrix alignment.

### EdU and immunostaining

Approximately 8-10 μL of 10 mM EdU solution were injected intraperitoneally (IP) into adult zebrafish 24 hours prior to tissue collection. For whole mount EdU-staining, adult heads were fixed overnight in 4% paraformaldehyde at 4 degrees. The tendons were the dissected out of the heads and permeabilized overnight in 1% Triton-X/D-PBS. Click-it EdU staining was then performed according to the manufacturer’s instructions (Invitrogen) followed by Draq5 nuclear counterstaining and 2-photon imaging. For the combined lineage tracing and EdU staining on sections, adult tendons were dissected and fixed overnight in 4% paraformaldehyde at 4 degrees, brought up a gradient to 30% sucrose and embedded into OCT. Samples were cryo-sectioned at 12 μm thickness and blocked for 1 hour (8% donkey serum, 0.3% BSA, 1% Triton-X) prior to performing Click-it EdU staining. After EdU staining, immunostaining was performed with an anti-GFP antibody (1:250 dilution, Abcam Cat No. ab290) to stain for CFP/YFP labeled cells. Sections were then imaged using confocal microscopy.

### 2-photon imaging quantification

All imaging quantification was performed in a blinded manner. For both the immune cell and EdU quantification across timepoints, 2-3 slices were extracted from the z-stacks of the whole mounted tendon tissue from individual animals. The images were processed to include only the injury area (i.e. between the two severed tendon ends), blinded, and the percentage of either *mpx:eGFP*+, *mpeg:eGFP*+, or EdU+ cells out of total Draq5+ cells in the injury site was quantified. The percentages across all sections for each sample were averaged. A one-way ANOVA statistical analysis was performed with Tukey’s multiple comparisons tests to determine statistically significant changes across the dataset and between different timepoints.

### Generation and validation of a TgBAC(scxa:creERT2) zebrafish line

To generate a *TgBAC(scxa:creERT2)* zebrafish line, standard BAC recombineering was performed to modify the *scxa*-containing BAC (CH211-251G8) from the CHORI BAC library to 1) include a *cryaa:sfGFP* in the backbone of the BAC for screening purposes and 2) introduce a creERT2 immediately after the *scxa* promoter. Briefly, to insert *cryaa:sfGFP* into the backbone of the BAC, the iTol2 flanked kanR-cryaa:sfGFP fragment was PCR amplified from Addgene #74153 (Fuentes et al., 2016) and electroporated into *scxa*-containing BAC containing cells. A successfully recombineered clone was identified and utilized for subsequent steps. To introduce the creERT2 sequence after the *scxa* promoter, a custom vector housing a creERT2-frt-ampR-frt fragment was first generated via Gibson assembly. The creERT2-frt-ampR-frt fragment was then amplified with primers containing homology arms for the *scxa* locus such that the insertion should occur directly into the ATG start site. The fragment was then recombineered into the *scxa* BAC with cryaa:sfGFP to generate the final *scxa:creERT2* BAC construct.

Wildtype Tübingen zebrafish eggs were injected at the 1 cell stage with the engineered *scxa:creERT2* BAC and *tol2* mRNA. Injected F0 larvae that displayed green lenses were grown to adulthood for germline transmission screening and validation of creERT2 labeling. To screen for potential founders, F0 adults were outcrossed to the *ubi:zebrabow* line and embryos were immersed in 20 μM 4-hydroxytamoxifen (4OH-T) beginning at 48 hpf. Developing larvae were screened at 72 hpf to identify founders that gave rise to progeny with labeled developing *scxa+* tendon structures. Double *scxa:creERT2;ubi:zebrabow* embryos from promising founders were then raised to adulthood and dosed with 4OH-T to test for labeling in the adult MST. Of these, 3 stable founder lines were established all showing high specificity, but variable labeling efficiency during adulthood. Therefore, we moved forward with the founder line showing highest labeling efficiency for all subsequent experiments. To optimize labeling, several routes of 4OH-T delivery were tested including immersion, IP injection, and local injection. Of these, the highest efficiency of labeling was determined to be via immersion.

### Lineage tracing of adult scxa:creERT2 zebrafish

Adult zebrafish (~5-6 months old) were immersed in 2.5 μM 4OH-T for 3 nights (~12 hours total) at a density of 100 mL per fish and protected from light. Between treatments, fish were placed into normal fish system water and fed daily. Following the conclusion of 4OH-T treatment, fish were returned to the normal system and tendon injuries were performed at 7 or 14 days post-treatment depending on the experiment. Tissue was collected at designated timepoints and either imaged on the 2-Photon or processed for combined EdU/antibody staining (see prior section).

## Author Contributions

S.L.T., R.R.S., and J.L.G. conceptualized the study design. S.L.T. performed all the experiments, data analysis, and wrote the manuscript. S.V. assisted in generating the *Tg(scxa:creERT2)* line, TEM analysis, and image quantification. S.V., R.R.S., and J.L.G. revised the manuscript. All authors approved the final manuscript.

## Acknowledgements

We would like to thank Sara Tufa and Doug Keene at the Shriners Hospital for Children in Portland, Oregon for their help with performing the TEM. We would also like to thank our funding sources for this work: S.L.T. was supported by the Helen Hay Whitney Foundation (HHWF) Postdoctoral Fellowship. S.V., R.S., and J.L.G. were supported by the Harvard Stem Cell Institute, NIH/NIAMS AR074541 and AR079495. The funding sources played no role in study design, data collection, analysis and interpretation of data, or the writing of this manuscript.

## Data Availability

The authors declare that all the data generated in this study are available in the manuscript and supplemental information.

