## Supplemental Materials for "Endogenous Tenocyte Activation Underlies the Regenerative Capacity of Adult Zebrafish Tendon"

### Supplemental figures and legends

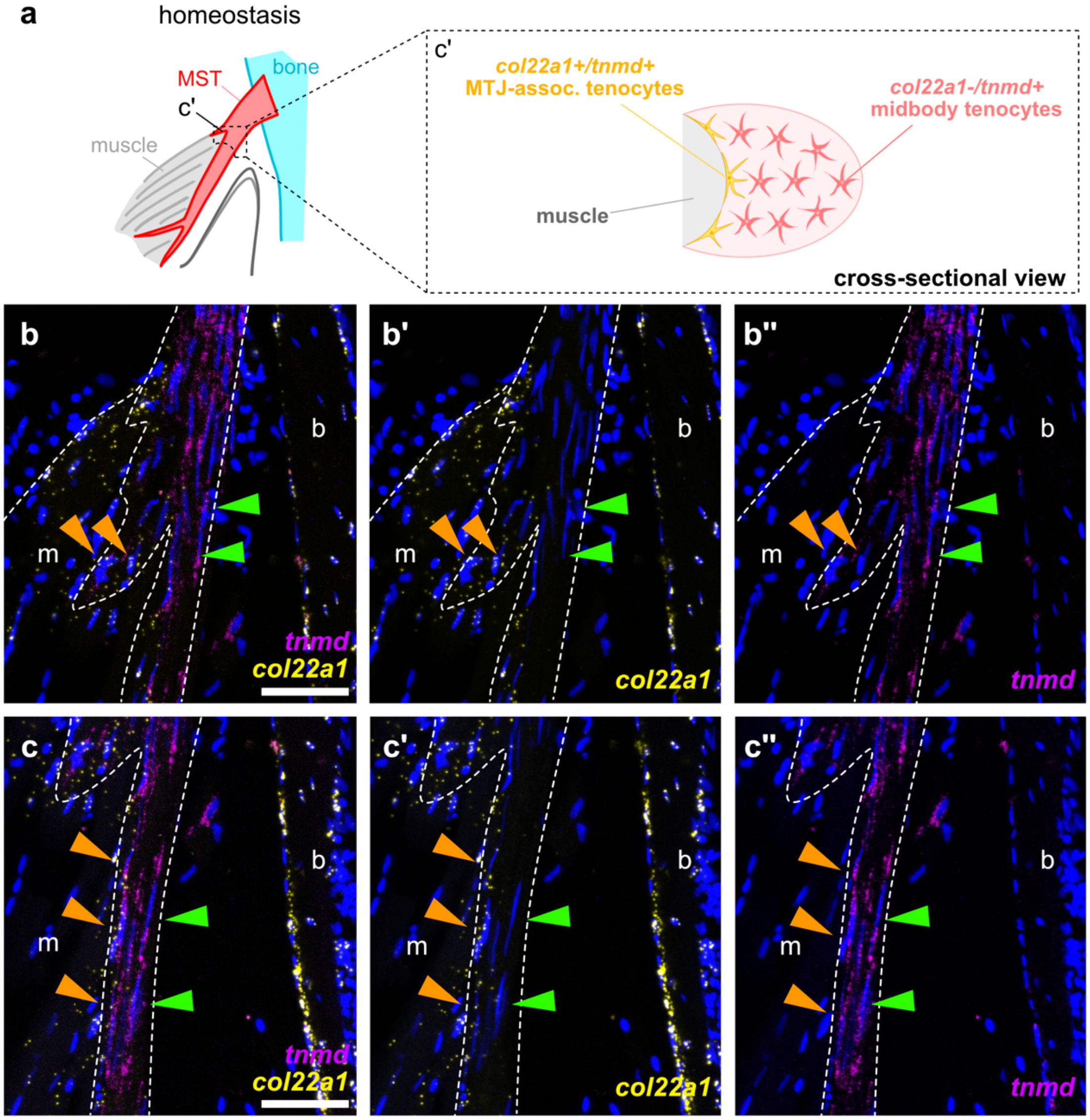


**Supplemental Figure 1. The MST contains cells expressing midbody tenocyte and myotendinous junction markers.**

(a) Graphical schematic showing a cross-sectional view of the MST during homeostasis to depict the cellular composition based on the expression patterns shown in c'.

(b-c'') Section RNAscope *in situ* hybridization of *tnmd* and *col22a1* on an uninjured adult control MST showing expression along two different regions of the tendon. Panels b-b'' show a region at the beginning of the MTJ and the midbody/MTJ section. Panels c-c'' show the section of the midbody that contains MTJ cells along the length of the tendon. *Col22a1* expression is specifically localized to the tendon cells that are in direct contact (*tnmd+/col22a1+*) with the muscle (m) (orange arrowheads), while *tnmd+/col22a1-* cells are present in tenocytes not directly in contact with the muscle along the length of the tendon (green arrowheads). *Col22a1+* cells can also be seen in periosteal cells lining the maxillary bone (b).

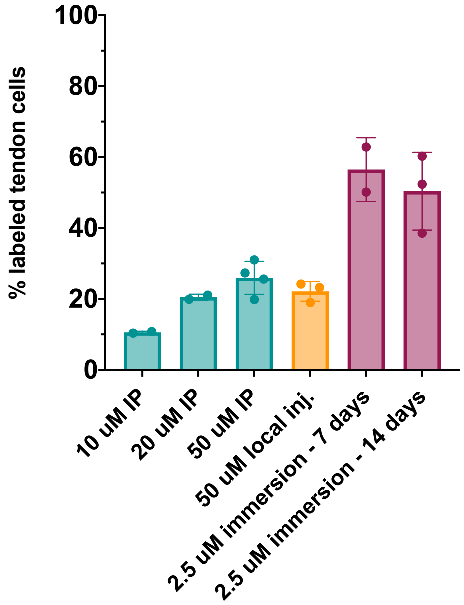


**Supplemental Figure 2. Optimizing 4OH-T delivery to obtain maximal tendon labeling during homeostasis.** Quantification of the percentage of CFP+ and/or YFP+ labeling in *scxa:creERT2;ubi:zebrabow* zebrafish after various methods of 4OH-T delivery. For the IP injections, ~10 µL of 4OH-T at the various concentrations detailed was injected daily for 3 consecutive days. For the local injection, ~10 µL 50 µM of 4OH-T was injected locally into the cavity between the skin and the jaw adductor muscle attached to the MST. For immersion, zebrafish were incubated for 3 nights in 2.5 µM of 4OH-T. For all conditions, the labeling was analyzed 7 days after the last treatment, except for immersion which was assessed at both 7 and 14 days post-induction. IP, intraperitoneal; dps, days post-induction.
